# Citrullination of proteins as a specific response mechanism in plants

**DOI:** 10.1101/2020.09.12.294728

**Authors:** Claudius Marondedze, Giuliano Elia, Ludivine Thomas, Aloysius Wong, Chris Gehring

## Abstract

Arginine deamination, also referred to as citrullination of proteins by peptidyl-arginine deiminases, is a post-translational modification affecting histone modifications, epigenetic transcriptional regulation and proteolysis in animals, but has not been reported in higher plants. Here we report, firstly, that *Arabidopsis thaliana* proteome contains proteins with a specific citrullination signature and that many of the citrullinated proteins have nucleotide-binding regulatory functions. Secondly, we show that changes in the citrullinome occur in response to cold stress, and thirdly, we identify an *Arabidopsis thaliana* protein with calcium-dependent arginine deiminase activity. Taken together, these findings establish this post-translational modification as a hitherto neglected component of cellular reprogramming during stress responses.

## INTRODUCTION

Citrulline, an intermediate of the urea cycle, can be formed in proteins and peptides as a result of arginine deamination (Figure 1A). The presence of citrulline in protein extracts was first reported well over half a century ago in the red alga *Chondrus crispus* (Smith and Young, 1955). It was later discovered in mammalian hair follicles and established unequivocally in peptide linkages (Rogers, 1962; Rogers and Simmonds, 1958). Enzymatic conversion of an arginine to a citrulline in a protein results in the loss of a positive charge that can perturb intra- and inter-molecular electrostatic interactions causing partial unfolding of the citrullinated protein (Tarcsa et al., 1996). Disturbances of cellular citrullination signatures have been implicated in apoptosis, the regulation of pluripotency (Christophorou et al., 2014), cancer (Yuzhalin, 2019) and autoimmune and neurological disorders (Klareskog et al., 2013; van Venrooij and Pruijn, 2000). Citrullination also has a role in the epigenetic regulation of transcription (Thompson and Fast, 2006) and several types of histones (H2A, H3 and H4) have been shown to be citrullinated by the peptidyl arginine deiminase type IV (PAD4) in the nucleus of granulocytes where the reaction uses both arginine and methylarginine (but not dimethylarginine) as substrates. PADs have been discovered in several vertebrate species including mammals, but have not been observed in bacteria, lower eukaryotes or plants (György et al., 2006; Joshi and Fernie, 2017). However, citrullination as a post-translational modification catalyzed by PADs has not been reported in plants, on the contrary, it has been suggested that the modulation of protein function through citrullination was an unlikely proposition (for review see (Gudmann et al., 2015)).

**Figure 1.**
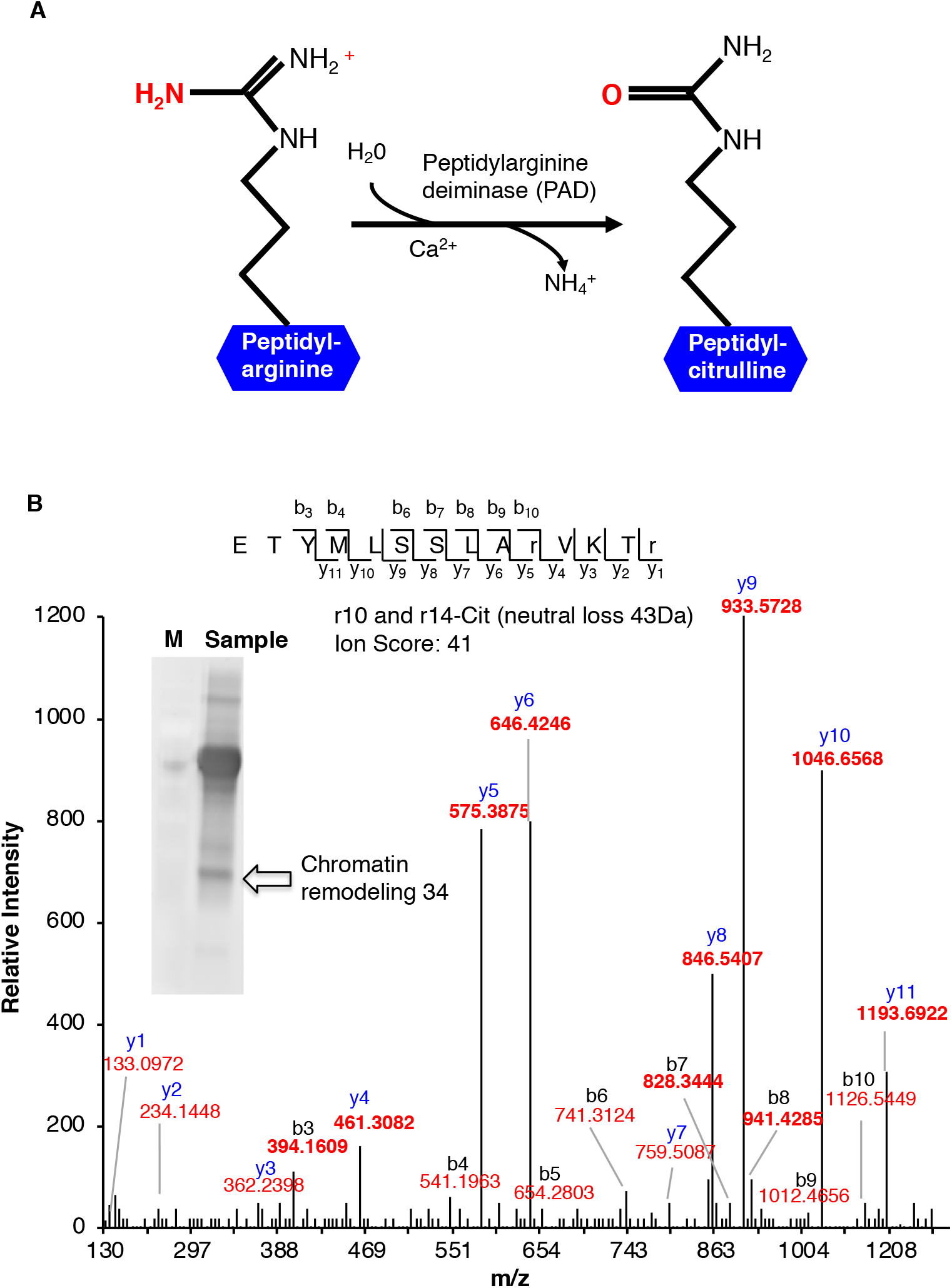
(A) Enzymatic formation of a citrulline. Enzymatic hydrolysis of peptidyl-arginine to peptidyl-citrulline by peptidylarginine deiminase (PAD). (B) MS/MS spectra of a citrullinated Chromatin Remodeling 34 (ChR34). The MS/MS spectrum of Chromatin Remodeling 34 (At2g21450) contains citrullinated residues on position 10 and 14. Indicated on the peptide sequence are the detected fragment ions, as indicated by the b- and y-ions and their relative intensities. Ion score of the peptide is 41. The insert shows a western blot of anti-citrulline immunoprecipitated proteins. The arrow shows the protein band identified as “Chromatin Remodeling 34”.

Here we address the following questions: (1) Is there evidence for citrullinated proteins in *Arabidopsis thaliana*, and if so, (2) are citrullination signatures stimulus-specific and (3), what are plausible *Arabidopsis thaliana* deiminase candidates that can citrullinate proteins.

## RESULTS AND DISCUSSION

### In search of citrullinated Arabidopsis thaliana proteins

The experimental system is *Arabidopsis thaliana* (ecotype Columbia-0) cell culture cells subjected to nuclear fraction enrichment and tested for citrullinated proteins by immunoprecipitation followed by western blotting with anti-citrullination antibodies (Supplemental Figure 1). The presence of candidate proteins was further validated by tandem mass spectrometry (LC-MS/MS) following in-blot digestion and/or direct in-solution digestion of the immunoreactant proteins and this revealed several citrullinated peptides (Table 1) and notably peptides from proteins with DNA or RNA targets indicating that this post-translational modification is likely to have a role in cellular (re-)programming during development and / or stress responses. A point in case are <u>Ch</u>romatin <u>R</u>emodelling 34 (ChR34; At2g21450; Figure 1B) with two out of 52 arginine residues citrullinated, and the RING/FYVE/PHD zinc finger protein (At3g02890), that are both strongly expressed in reproductive organs and distinct stages of seed formation (see https://genevestigator.com/gv/ (Zimmermann et al., 2004)).

**Table 1.** Citrullinated peptides and low temperature stress-induced citrullination events in *Arabidopsis thaliana* proteins.

| Accession number | Description | Mascot score | Mass | Peptides | Pep. Score (≥30) | Expect | FC C vs 1h | FC C vs 24h |
| --- | --- | --- | --- | --- | --- | --- | --- | --- |
| **Citrullinated peptides in the control** |  |  |  |  |  |  |  |  |
| At1g50030<sup>#</sup> | Target of rapamycin (TOR) | 59 | 281237 | R.ERAVEALR.A | 59 | 0.00031 | ns | ns |
| At1g26110<sup>#</sup> | Decapping 5 (DCP5) | 47 | 64331 | R.GYGGyGGRGGGGGGYGYGGRGQGR.G | 47 | 0.007 | ns | ns |
| At5g17860<sup>#</sup> | Calcium exchanger 7 (CAX7) | 40 | 62679 | K.RMSDQILR.S | 39 | 0.033 | ns | ns |
| At4g12850<sup>#</sup> | Far-red impaired responsive (FAR1) | 37 | 21522 | K.VRDIVENVK.L | 36 | 0.035 | ns | ns |
| At4g31890<sup>#</sup> | Armadillo (ARM) repeat protein | 35 | 56650 | R.VtLAMLGAIPPLVSmIDDsR.I | 35 | 0.0097 | ns | ns |
| At4g27810<sup>#</sup> | Hypothetical DUF688 protein | 34 | 22156 | R.RSLSVIR.R | 34 | 0.031 | ns | ns |
| At3g02890<sup>#</sup> | RING/FYVE/PHD zinc finger | 33 | 110280 | R.RVGNRPMGRR.G | 33 | 0.022 | ns | ns |
| At4g00830<sup>#</sup> | RNA-binding family protein | 33 | 55273 | R.NNGSSGGSGRDNSEHDGNGRGGR.R | 33 | 0.027 | ns | ns |
| At4g09980<sup>#</sup> | Methyltransferase MT-A70 | 32 | 86022 | R.ERTHGSSDSSK.R | 32 | 0.024 | ns | ns |
| At4g10000<sup>#</sup> | Thioredoxin family protein | 30 | 37375 | R.ISGNGNWVRER.R | 30 | 0.018 | ns | ns |
| **Peptides differentially citrullinated in response to cold treatment** |  |  |  |  |  |  |  |  |
| At2g21450 | Chromatin remodeling 34 (ChR34) | 68 | 94037 | K.ETYmLSSLARVKTR.R | 41 | 0.0018 | 1.90 | 1.84 |
| At5g27920<sup>#</sup> | F-box family protein | 41 | 72636 | R.ARGLETLMAR.M | 47 | 0.0029 | 1.86 | 1.53 |
| At4g19180 | GDA1/CD39 nucleoside phosphatase | 37 | 82544 | R.FQRWSPMSTGVK.T | 37 | 0.0077 | 1.9 | 1.8 |
| At1g29040<sup>#</sup> | Hypothetical exostosin family prot. | 33 | 20111 | K.FLSSRSK.Q | 33 | 0.013 | ns | 0.67 |
| **De novo cold-induced citrullinated peptides** |  |  |  |  |  |  |  |  |
| At1g24290<sup>#</sup> | AAA-type ATPase family protein | 37 | 57702 | K.SMRGGDANAAYWLAR.M | 35 | 0.013 | at 1h | at 24h |
| ATC00520 | YCF4 (unfolded protein binding) | 35 | 21564 | K.DIQsRIEVK.E | 35 | 0.014 | at 1h | nd |
| At1g16950 | Hypothetical protein | 34 | 10044 | .MARPRIsImICLLILIVGFVLQssQAR.K | 34 | 0.0071 | nd | at 24h |
| At2g23790 | Unknown DUF607 protein | 32 | 38438 | K.LLRAAQIEIVK.S | 32 | 0.02 | at 1h | nd |
| At2g47650 | UDP-xylose synthase 4 | 32 | 50085 | R.VVvtGGAGFVGSHLVDRLmAR.G | 32 | 0.0087 | nd | at 24h |

Citrullinated residues in peptides are in red; # Identified by both gel-based and gel-free techniques; ns: not significant at $p \leq 0.05$ and $FC = 0.68 < x < 1.5$ ; nd - not identified.

In order to detect possible stimulus-specific changes in citrullination signatures, we decided to test the effects of low temperature exposure of cells since it can be easily and reproducibly applied to suspension culture cells. Four proteins showed differential expression and degree of citrullination over time (0, 1 h and 24 h) (Table 1). They include the Chromatin Remodelling 34 (ChR34; At2g21450) (see spectrum in Figure 1B), an F-box family protein (At5g27920), GDA1/CD39 nucleoside phosphatase (At4g19180) and hypothetical exostosin family protein (At2g29040). It is noteworthy that these proteins are all expressed in the male gametophyte, which is most sensitive to cold (Dekkers et al., 2013). Five proteins undergo *de novo* citrullination of unmodified arginines in response to cold stress (Table 1). In the AAA-type ATPase (At1g24290), annotated as having a role in DNA replication, citrullination is observed at 1 h and 24 h after the onset of cold stress. Other proteins display different temporal signatures, e.g. the YCF4 chloroplast protein (AtCg00250) which has a role in translation and DNA-templated elongation, is citrullinated only at 1h.

### Identification of a candidate arabidopsis peptidyl-arginine deiminases

In order to find candidate arabidopsis peptidyl-arginine deiminases, also referred to as agmatine deiminases, we aligned key amino acid residues in the catalytic center of annotated bacterial peptidyl-arginine deiminases and created a search motif: W-X-R-D-[TS]-G-X(100,140)-H-[VIL]-D (Figure 2A) and identified At5g08170. Based on the crystal structure of At5g08170 (PDB ID: 3H7K), the key amino acids appearing in the motif were located and the conserved amino acids constitute a clear cavity (highlighted in yellow) that is solvent exposed (Figure 2B) and can spatially accommodate an arginine residue (Figure 2B and 2C). Next, we modelled if arginines in two selected citrullinated proteins (At2g21450 and At4g00830) can dock at the assigned catalytic center of At5g08170. In order to simulate docking, models of both At2g21450 and At4g00830 citrullinated proteins were first generated by homolog modeling against the chain K of an ATPase domain of a chromatin remodeling factor (PDB ID: 6PWF) and the chain D of decaheme c-type cytochrome (PDB ID: 6R2K) respectively. Based on the 3D models, the citrullinated arginines (colored according to surface charges) were determined to be solvent exposed and protruding into space that is visibly large enough for the arginines to have unimpeded interactions with the catalytic center of At5g08170 (Supplemental Figure 2A and B). The individual citrullinated residues: R427 and R431 of At2g21450, and R477 and R487 of At4g00830, were docked at the catalytic center cavity of At5g08170, keeping all bonds in the R ligand non-rotatable so that their poses from the generated 3D models, are preserved. All citrullinated arginines docked at the catalytic cavity in a binding pose deemed suitable for catalysis i.e., with the amine rich region pointing into the cavity, as shown in the surface models (right panels) (Supplemental Figure 2A and B).

**Figure 2.**
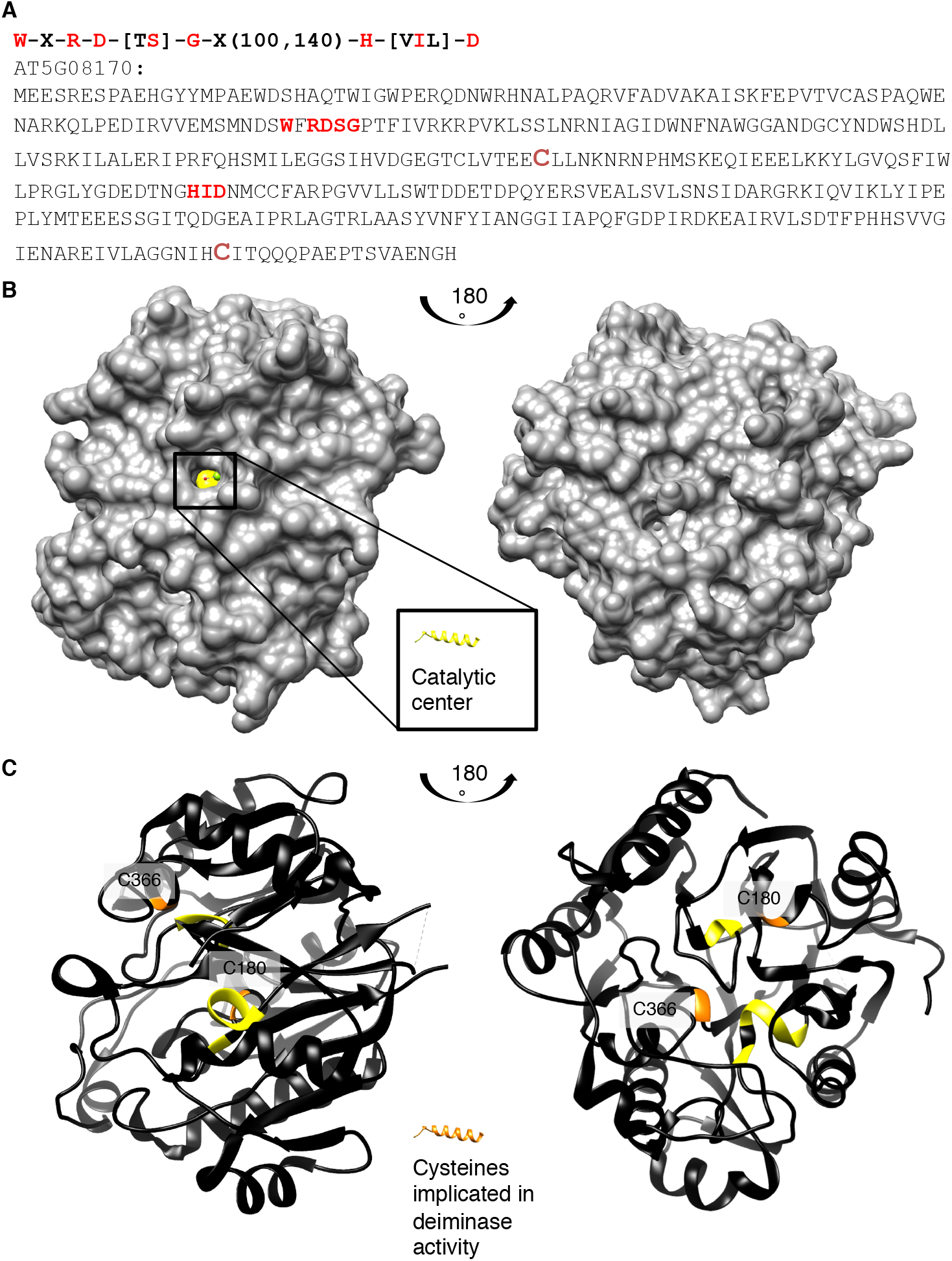
(A) Amino acid sequence, (B) surface and (C) ribbon structure of At5g08170 (PDB ID: 3H7K). Amino acids appearing in the motif: W-X-R-D-[TS]-G-X(100,140)-H-[VIL]-D are highlighted in red in (A) the full-length At5g08170 amino acid sequence, and colored yellow in (B) the surface and (C) ribbon At5g08170 structures. The cysteines (C180 and C366) implicated in catalytic activity of agmatine deiminase are highlighted in green in (a) the amino acid sequence and orange in (B) the surface and (C) the ribbon structures. Amino acids R93 – G96 from the motif, occupy a clear cavity that can spatially accommodate an R residue (B), thus is assigned the catalytic center for arginine deiminase (highlighted in yellow).

### Testing of peptidyl-arginine deiminase activity

*In vitro* citrullination with recombinant peptidyl-arginine deiminases was experimentally tested with various substrates including fibrinogen (Sigma-Aldrich, St. Louis, MO) and peptides generated from the “Like heterochromatin protein 1-Interacting Factor 2” (LIF2) and ChR34. Fibrinogen has been shown to be a target of citrullination by peptidyl-arginine deiminases from *Porphyromonas gingivalis* (Lundberg et al., 2010) and is citrullinated by the recombinant peptidyl-arginine deiminase (Supplemental Table 1). Furthermore, we show that the recombinant enzyme can also citrullinate LIF2 (Supplemental Table 1) as well as itself (Supplemental Table 2). The observed Ca^2+^-dependent auto-citrullination of a peptidyl-arginine deiminase is entirely consistent with findings in mammalian orthologs (Andrade et al., 2010).

In conclusion, citrullination events do occur in higher plants, as shown in the case of *Arabidopsis thaliana* proteome. In addition, low temperature stress causes stimulus-specific protein citrullination of a set of proteins many of which have annotated roles in stress responses. Furthermore, our data also suggest that citrullination may be part of the general regulation of pluripotency. Finally, we also predict that specific citrullination could occur in response to biotic and abiotic stimuli and that stress specific citrullination signatures and their down-stream effects influence plant growth, development and stress responses.

## METHODS

### Arabidopsis cell suspension culture growth

Cells derived from roots of *Arabidopsis thaliana* (ecotype Columbia-0) were grown in 100 mL of Gamborg’s B-5 (Gamborg and Eveleigh, 1968) basal salt mixture (Sigma-Aldrich, St Louis, MO, USA) with 2,4-dichlorophenoxyacetic acid (0.5 µg mL^-1^) and kinetin (0.05 µg mL^-1^) in 250 mL sterile flasks in a growth chamber (Innova^®^ 43, New Brunswick Scientific Co., NJ, USA) at 120 rpm, with photosynthetic light set for 12 h light/12 h dark cycles at 23°C, and sub-cultured every seven days (Marondedze et al., 2015; Piazza et al., 2015; Turek et al., 2014). Cells were cold (4°C) treated and collected at 0, 1 hour and 24 h post-treatment. Media was drained off using Stericup^®^ filter unit (Millipore, Billerica, MA), and cells were immediately frozen in liquid nitrogen and stored at -140°C until further use.

### Nuclear protein extraction

Nucleus enrichment was performed using a nucleus enrichment kit for tissue (Pierce, Rockford, IL, USA). Briefly, protease inhibitors were added to the appropriate volume of nucleus enrichment reagents A and B. A total of 500 mg of frozen callus cells was transferred to a 50 mL Falcon tube and 800 µL of reagent A and 800 µL of reagent B were added. The tissue was homogenized for 4 sec twice on ice using a PowerGen 125 grinder (Fisher Scientific, Rockford, IL, USA). The homogenate was transferred to a fresh 2 mL microcentrifuge tube, mixed by inverting the tube five times and centrifuged at 500 × g for 10 min. at 4°C to collect the supernatant. A discontinuous density gradient was prepared by carefully overlaying three OptiPrep™ gradients (first layer consisted of 1.5 mL of 27.5% (v/v), the second of 1.5 mL of 23% (v/v) and the third of 1 mL of 20% (v/v)). Gradients were prepared from the OptiPrep media and gradient dilution buffer. The tissue extract was mixed with 60% (v/v) OptiPrep™ cell separation media and diluted to a final density gradient of 7.5% (v/v) with gradient dilution buffer and overlaid on top of the density gradients. Ultracentrifugation was performed at 40,000 × *g* for 90 min. at 4°C. The top 2 mL, containing the nuclei, was pipetted out and precipitated with ice-cold acetone overnight at −20°C. Proteins were pelleted at 3,900 × *g* for 15 min. at 4°C, washed three times in 80% (v/v) ice-cold acetone, solubilized in urea lysis buffer (7 M urea, 2 M thiourea) as described in (Marondedze et al., 2015). Protein concentration was estimated by Bradford assay (Bradford, 1976). Approximately 200 µg of total nucleus protein extract was reduced with 5 mM dithiothreitol, alkylated, and used for protein digestion with sequencing-grade modified trypsin (Promega, Madison, WI, USA). Resulting peptides were purified using Sep-Pak Vac tC18 100 mg cartridge (Waters, Milford, MA, USA), as described previously (Groen et al., 2013) completely dried in a Speed Vac concentrator (Thermo Scientific, Bremen, Germany) and analysed by tandem mass spectrometry (LC-MS/MS).

### Immunoprecipitation, western blot and quantitative analysis of citrullinated proteins

Immunoprecipitation was performed using two protocols. In the first experiment immunoprecipitation was done using immunoprecipitation kit Dynabeads^®^ Protein A (Life technologies) and according to manufacturer’s instruction. Here, Dynabeads^®^ were resuspended by vortexing >30 sec and 50 µL of Dynabeads^®^ were pipetted to a 1.5 mL microcentrifuge tube. The supernatant was removed using a magnetic Eppendorf tube rack prior to adding 2 µg of anti-citrulline antibody (ACAb, Abcam, Cambridge, UK) diluted in 200 µL phosphate buffered saline (PBS) containing Tween-20. The beads-Ab mix was incubated with rotation for 10 min. at room temperature. The supernatant was removed by placing the tube on the magnetic rack and the beads-Ab complex was resuspended in 200 µL PBS/Tween-20 buffer, resuspended by gentle pipetting and the supernatant was removed as stated previously. The protein sample (200 µg) was then added to the beads-Ab and gently pipetted to mix, incubated with rotation for 20 min. at room temperature and supernatant decanted. Beads-Ab-protein mix was washed three times using 200 µL of washing buffer for each wash and resuspended in 100 µL washing buffer by gentle pipetting and transferred to a fresh tube. Supernatant was removed by means of a magnetic rack. A 20 µL of elution buffer was gently pipetted to resuspend the Dynabeads^®^-Ab-protein complex and incubated with rotation for 2 min. at room temperature to dissociate the complex.

In the second experiment, 200 µg of protein sample and 2 µL of anti-citrulline antibody in TBS were mixed and incubated overnight at 4°C with gentle rocking. Dynabeads^®^ Protein A (Novex^®^, Life technologies) were then added and incubated for 3 h at 4°C with gentle rocking. The protein-antibody-beads complex was washed five times with 1 × lysis buffer diluted in TBS, resuspended in 20 µL of elution buffer, mixed gently and incubated with end-over-end mixing for 2 min. at room temperature to dissociate the complex. Supernatant was collected using a magnetic rack.

For both experiments, the protein solution collected was split into two-10 µL aliquots, one was used for in-solution digestion and tandem mass spectrometry and the remaining 10 µL was mixed with equal volume of 2 × reducing buffer for preparation prior to one-dimensional gel electrophoresis and western blot analysis. Proteins were transferred from polyacrylamide gel to polyvinylidine difluoride (PVDF) membranes as described by (Towbin et al., 1979) using a Trans-blot^®^ electrophoretic transfer cell (Bio-Rad, city, country). The PVDF membrane with transferred proteins was blocked overnight at 4°C in blocking solution (5% (w/v) bovine serum albumin in TBS), washed three times with TBS buffer containing 0.05% (v/v) Tween 20 (TBST) for 5 min. and incubated with anti-citrulline (primary antibody) diluted in TBST for 1 h at 37°C. Membranes were washed three times in PBST for 5 min., incubated with the secondary antibody, Alexa Fluor^®^ 488 goat anti-rabbit IgG (Molecular Probes, Eugene, OR, USA) diluted in TBS for 2 h at room temperature, thoroughly washed in TBST and imaged with a Typhoon™ 9410 scanner (GE Healthcare, city, country). Quantitative analyses and the relative-fold changes for protein bands were computed using ImageQuant-TL 7.0 software (GE Healthcare). Membranes were stained using the MemCode™reversible stain according to the manufacturer’s instructions (Pierce, city, country). All visibly stained bands were cut, digested with trypsin and analyzed by LC-MS/MS.

### Tandem mass spectrometry and database search

Peptides were analyzed by mass spectrometry as described previously (Marondedze et al., 2016; Marondedze et al., 2013). Briefly, peptides were re-suspended in 5% (v/v) acetonitrile and 0.1% (v/v) formic acid prior to identification by LTQ Orbitrap Velos mass spectrometer (Thermo Electron, Bremen, Germany) coupled with an Ultimate 3000 UPLC (Dionex-Thermo-Scientific) for nano-LC-MS/MS analyses. A volume of 4 µL of peptide mixtures was injected onto a 5 mm long × 300 µm C18 PepMap 100 µ-precolumn, 5 µm, 100 Å (Thermo-Scientific) and then to a 150 mm × 75 µm Acclaim PepMap 100 NanoViper C18, 3 µm, 100 Å (Thermo-scientific) column. The MS scan range was *m/z* 350 to 1600 and spray voltage 1500 V. Top 10 precursor ions were selected in the MS scan by Orbitrap with a resolution r = 60,000 for fragmentation in the linear ion trap using collision induced dissociation. Normalized collision-induced dissociation was set at 35.0. Spectra were submitted to a local MASCOT (Matrix Science, London, UK) server and searched against arabidopsis in the TAIR (release 10), with a precursor mass tolerance of 10 ppm, a fragment ion mass tolerance of 0.5 Da, and strict trypsin specificity allowing up to one missed cleavage, carbamidomethyl modification on cysteine residues as fixed modification and oxidation of methionine residues, citrullination (also known as deamination) of arginine residues as variable modifications. Proteins were considered positively identified if the Mascot score was over the 95% confidence limit corresponding to a score ≤32 and peptide was considered citrullinated if the Mascot ion score was ≤30.

### Structural analysis

Amino acid consensus motifs of catalytic centers were obtained as first employed in the search for plant nucleotide cyclases (Ludidi and Gehring, 2003) and the motif searches of Swiss-Prot were done on-line (https://www.genome.jp/tools/motif/MOTIF2.html). The crystal structure of *Arabidopsis thaliana* agmatine deiminase At5g08170 was obtained from protein data bank (PDB ID: 3H7K) and the location of the arginine citrullination catalytic center was identified and visualized using UCSF Chimera (ver. 1.10.1) (Pettersen et al., 2004). Two selected citrullinated proteins At2g21450 and At4g00830 were modeled against the chain K of a ATPase domain of a chromatin remodeling factor (PDB ID: 6PWF) and the chain D of decaheme c-type cytochrome (PDB ID: 6R2K) respectively using the Modeller (ver. 9.14) software (Sali and Blundell, 1993). The citrullinated arginines in the generated models were visualized and assessed for their ability to spatially fit the catalytic center of At5g08170. Docking simulations of the citrullinated arginines from the two peptides to the catalytic center of At5g08170 was performed using AutoDock Vina (ver. 1.1.2) (Sali and Blundell, 1993; Trott and Olson, 2010). All bonds in the ariginines were set as non-rotatable to keep their poses similar to that in the peptides. The arginine docking poses were analyzed, and all images created by UCSF Chimera (ver. 1.10.1) (Pettersen et al., 2004). Chimera was developed by the Resource for Biocomputing, Visualization, and Informatics at the University of California, San Francisco (supported by NIGMS P41-GM103311).

### Cloning and Expression of At5g08170

Total RNA from arabidopsis (Col-0) leaves was extracted using RNeasy Plant Mini Kit (Qiagen). Followed by synthesis of cDNA using SuperScript™ II Reverse Transcriptase system (Invitrogen). Full length At5g08170 was then detected and amplified by polymerase chain reaction (PCR) using the forward primer (5’-ATGGAGGAGTCACGAGAATCG-3’) and reverse primer (5’-TCAGTGGCCATTTTCGGC-3’). PCR amplified gene was cloned into pCR8™/GW/TOPO^®^ plasmid prior to clone it into pDEST™ 17 destination vector using GATWAY cloning technology (Invitrogen). Plasmid contained the full length gene was then transformed into *E. coli* BL-21 A1 cells (Invitrogen). The expression of recombinant protein was induced by 0.2% (w/v) L-Arabinose (Sigma Aldrich). The expressed protein was purified as His-tagged fusion protein using Ni-NTA agarose beads (Qiagen) and analyzed by SDS-PAGE.

### Enzyme activity analysis via mass spectrometry

Two synthetic peptides from two of the identified citrullinated proteins, LIF2 and chromatin remodelling 34 were designed and purchased (Thermo Fisher Scientific). The selected peptides were designed to ensure that all peptides had at least one arginine or lysine to confirm the success of digestion. A tryptic digestion was performed following a citrullination assay that was performed in 50 mM CHES, pH 9.5, 5mM CaCl_2_, 10 mM DTT, 50 µg of different substrates (including CHES buffer as control, the two synthetic peptides and fibrinogen) and 50 µg of purified At5g01870 in a total volume of 200µl in microcentrifuge tube. Notably, autocitrullination of the agmatine deiminase in the presence and absence of calcium was also assessed by tandem mass spectrometry as described earlier.

## FUNDING

G.E. was the recipient of an Erasmus Mundus Action 2, Strand 2, lot 4 (Gulf Countries) award. A.W. was supported by the National Natural Science Foundation of China (31850410470) and the Zhejiang Provincial Natural Science Foundation of China (LQ19C130001).

## AUTHOR CONTRIBUTIONS

G.E. and C.G. conceived of the project, C.M. and C.G. planned the experiments, C.M. and L.T. did the experiments and A.W. did the structural modeling. All authors contributed to the writing of the manuscript.

## ACKNOWLEDGMENTS

We thank Lee Staff for carefully reading the manuscript. No conflicts of interest are declared.

## Supplemental Information (SI)

**Supplemental Figure 1:**
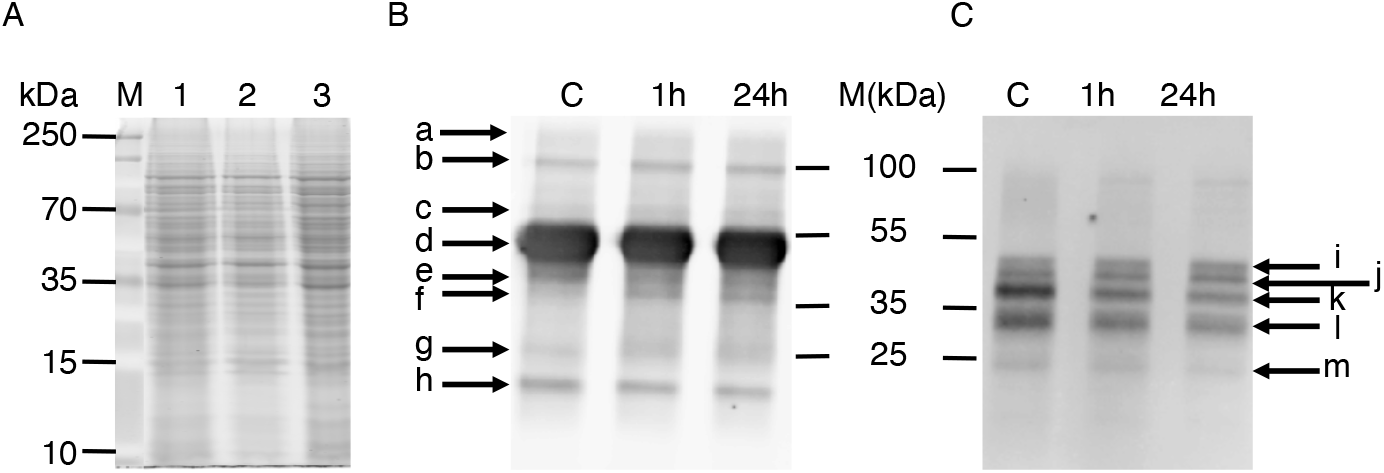
One-dimensional gel electrophoresis scans showing nuclear enriched proteins (A), western blots of immunoprecipitated citrullinated candidate proteins (B and C). In panel A, lane 1 represent the protein ladder (M), lane 2 -nuclear extract from the control samples of Arabidopsis cell suspension culture, lane 3 -nuclear extract from 1 hour cold treated Arabidopsis cell suspension culture sample and lane 4 -nuclear extract from 24 hour cold treated Arabidopsis cell suspension culture sample. For the Western blots the three lanes represent control (C), I hour post cold treatment (1h) and 24 hour post cold treatment (24h). Panel B represents a western blot where the nuclear extract was incubated overnight with the anticitrulline antibody and then conjugated with IgA protein beads. Panel C represents a western blot where the anticitrulline antibody was conjugated with IgA protein beads and then incubated with the nuclear extract for 10 minutes according to the manufacturer’s instruction (see supplementary document, Materials and Methods). In Panel B; **a**=AAA-type ATPase family protein, target of rapamycin, methyltransferase A70, far-red impaired responsive, calcium exchanger 7; **b**=chromatin remodeling 34, unknown protein, ARM repeat superfamily protein; **c**= no positive identification (ND); **d**=RNA-binding (RRM/RBD/RNP motifs) family protein; **e**=Unknown protein; **f**=AAA-type ATPase family protein; **g**=chromatin remodeling 34, GDA1/CD39 nucleoside phosphatase; **h**=RING/FYVE/PHD zinc finger superfamily protein. In panel C; **i**=F-box family protein; **j**=Decapping 5, thioredoxin family protein; **k**=AAA-type ATPase family protein, RNA-binding (RRM/RBD/RNP motifs) family protein, **l**=peroxidase superfamily protein; **m**= chromatin remodeling 34.

**Supplemental Figure 2:**
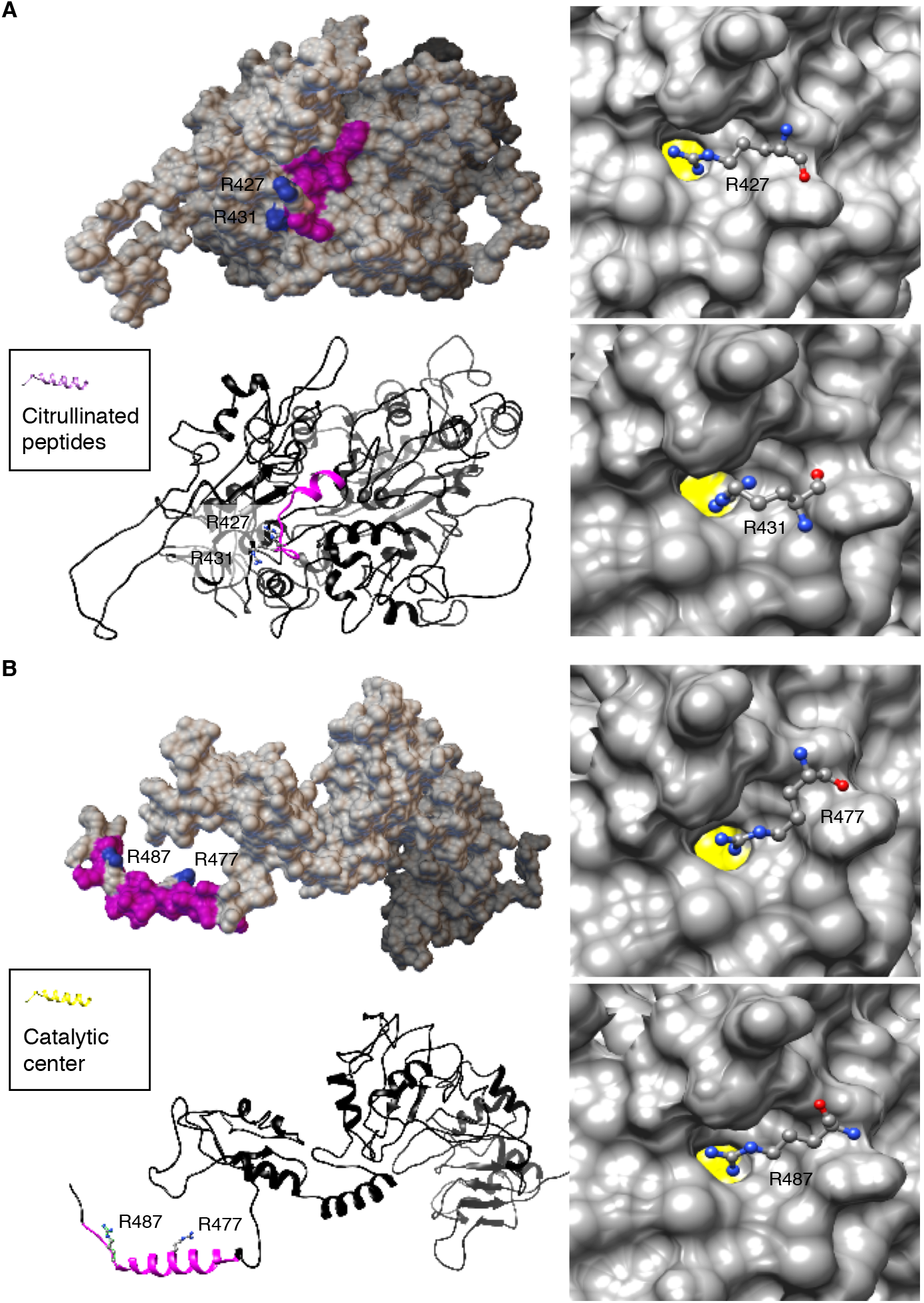
Computational assessment of the citrullinated arginines of two selected proteins (A) At2g21450 and (B) At4g00830. At2g21450 and At4g00830 were modeled against the chain K of a ATPase domain of a chromatin remodeling factor (PDB ID: 6PWF) and the chain D of decaheme c-type cytochrome (PDB ID: 6R2K) respectively using the Modeller (ver. 9.14) software. The citrullinated arginines (colored according to surface charges) in the generated models were visualized and assessed for their ability to spatially fit the catalytic center of At5g08170. Citrullinated peptides were colored magenta and citrullinated arginine residues are all solvent exposed as shown in the ribbon and surface models of At2g21450 and At4g00830 respectively (left panels). Individual citrullinated residue: R427 and R431 of At2g21450, and R477 and R487 of At4g00830, were respectively docked at the catalytic center cavity of At5g08170, keeping all bonds in the R ligand non-rotatable so that their poses in the generated 3D models are retained. All citrullinated arginines docked at the catalytic cavity in a binding pose deemed suitable for catalysis i.e., with the amine rich region pointing into the cavity, as shown in the surface models (right panels). All docking simulations were performed using AutoDock Vina (ver. 1.1.2) [48]. Docking poses were analyzed, and all images created using UCSF Chimera (ver. 1.10.1). Chimera was developed by the Resource for Biocomputing, Visualization, and Informatics at the University of California, San Francisco (supported by NIGMS P41-GM103311).

**Table S1.** Supplemental Table 1: Citrullinated fibrinogen and LHP1-INTERACTING FACTOR 2 RNA binding protein peptides following incubation with agmatine deiminase

| **A. Commonly citrullinated peptides following citrullination of fibrinogen with plant agmatine deiminase** |  |  |  |
| --- | --- | --- | --- |
| **Protein accession** | **Protein description** | **Peptide sequence** | **Literature\*** |
| gi\|237823914 | chain A, Human fibrinogen | ADSGEGDFLAEGGGVrGPR | X |
| gi\|182439 | fibrinogen gamma chain | ANQQFLVYCEIDGSGNGWTVFQKr | X |
| gi\|237823915 | chain B, Human fibrinogen | EEAPSLrPAPPPISGGGYR | X |
| gi\|4503689 | fibrinogen α-E preproprotein | GDFSSANNrDNTYNR | X |
| gi\|237823915 | chain B, Human fibrinogen | GGETSEMYLIQPDSSVKPYrVYCDMNTENGGWTVIQNr | X(C-Term.) |
| gi\|4503689 | fibrinogen α-E preproprotein | GGTSYGTGSETESPrNPSSAGSWNSGSSGPGSTGNr | X |
| gi\|4503689 | fibrinogen α-E preproprotein | GGTSYGTGSETESPrNPSSAGSWNSGSSGPGSTGNR | X |
| gi\|4503689 | fibrinogen α-E preproprotein | HrHPDEAAFFDTASTGK | X |
| gi\|237823916 | chain C, Human fibrinogen | IHLISTQSAIPYALrVELEDWNGR |  |
| gi\|182439 | fibrinogen gamma chain | IHLISTQSAIPYALrVELEDWNGR |  |
| gi\|182439 | fibrinogen gamma chain | IIPFNrLTIGEGQQHHLGGAK | X |
| gi\|4503689 | fibrinogen α-E preproprotein | QFTSSTSYNrGDSTFESK | X |
| gi\|4503689 | fibrinogen α-E preproprotein | TFPGFFSPMLGEFVSETESrGSESGIFTNTK | X |
| gi\|182439 | fibrinogen gamma chain | VELEDWNGrTSTADYAMFK | X |
| gi\|237823915 | chain B, Human fibrinogen | VYCDMNTENGGWTVIQNrQDGSVDFGR | X |
| gi\|237823915 | chain B, Human fibrinogen | VYCDMNTENGGWTVIQNrQDGSVDFGr |  |
| **B. LHP1-INTERACTING FACTOR 2 RNA binding protein peptides that are citrullinated** |  |  |  |
| **Protein accession** | **Protein description** | **Peptide sequence** | **Peptide score** |
| AT4G00830.1 | RNA-binding protein | NDrNNGSSGGSGRDNSHEHDGNR | 50 |
| AT4G00830.1 | RNA-binding protein | NDrNNGSSGGSGrDNSHEHDGNR | 50 |

\*- Fibrinogen peptides citrullinated by rmPAD2 (van Beers et al., 2010)

**Table S2.** Supplemental Table 2: Auto-citrullination of agmatine deiminase in the presence or absence of calcium

| Agmatine peptide | Citrullinated site(s) | Stoichiometry (%; A,B) |
| --- | --- | --- |
| ESPAEHGYMPAEWDSHAQTWIGWPErQDNWr | 32, 37 | 13, 25 |
| ESPAEHGYMPAEWDSHAQTWIGWPErQDNWR | 32 | 30, 37 |
| FEPVTVCASPAQWENArK | 73 | 11, 17 |
| GLYGDEDTNGHIDNMCCFArPGVVLLSWTDDETDPQYEr | 233, 252 | 14, 18 |
| GLYGDEDTNGHIDNMCCFArPGVVLLSWTDDETDPQYER | 233 | 49, 35 |
| LAASYVNFYIANGGIIAPQFGDPIrDK | 331 | 20, 43 |
| LYIPEPLYMTEEESSGITQDGEAIPrLAGTr | 301, 306 | 11, 33 |
| LYIPEPLYMTEEESSGITQDGEAIPrLAGTR | 301 | 33 |
| NIAGIDWNFNWGGANDGCYNDWSDLLVsrK | 144 | 10, 18 |
| QDNWrHNALPAQR | 37 | 14, 19 |
| QLPEDIrVVEMSMNDSWFR | 81 | 11, 2 |
| QLPEDIrVVEmsMNDSWFR | 81 | 2 |
| VVEMSMNDSWFrDSGPTFIVR | 93 | 44, 50 |

A: Agmatine - CaCl; B: Agmatine + CaCl

## Notes

### Competing Interest Statement

The authors have declared no competing interest.

## REFERENCES

Andrade, F., Darrah, E., Gucek, M., Cole, R.N., Rosen, A., and Zhu, X. (2010). Autocitrullination of human peptidyl arginine deiminase type 4 regulates protein citrullination during cell activation. Arthritis Rheum. 62, 1630–1640.

Bradford, M.M. (1976). A rapid and sensitive method for the quantitation of microgram quantities of protein utilizing the principle of protein-dye binding. Anal. Biochem. 72, 248–254.

Christophorou, M.A., Castelo-Branco, G., Halley-Stott, R.P., Oliveira, C.S., Loos, R., Radzisheuskaya, A., Mowen, K.A., Bertone, P., Silva, J.C., Zernicka-Goetz, M., et al. (2014). Citrullination regulates pluripotency and histone H1 binding to chromatin. Nature 507, 104–108.

Dekkers, B.J.W., Pearce, S., van Bolderen-Veldkamp, R.P., Marshall, A., Widera, P., Gilbert, J., Drost, H.-G., Bassel, G.W., Müller, K., King, J.R., et al. (2013). Transcriptional dynamics of two seed compartments with opposing roles in Arabidopsis seed germination. Plant Physiol. 163, 205–215.

Gamborg, O.L., and Eveleigh, D.E. (1968). Culture methods and detection of glucanases in suspension cultures of wheat and barley. Can. J. Biochem. 46, 417–421.

Groen, A., Thomas, L., Lilley, K., and Marondedze, C. (2013). Identification and quantitation of signal molecule-dependent protein phosphorylation. Methods Mol. Biol. 1016, 121–137.

Gudmann, N.S., Hansen, N.U., Jensen, A.C., Karsdal, M.A., and Siebuhr, A.S. (2015). Biological relevance of citrullinations: diagnostic, prognostic and therapeutic options. Autoimmunity 48, 73–79.

György, B., Tóth, E., Tarcsa, E., Falus, A., and Buzás, E.I. (2006). Citrullination: a posttranslational modification in health and disease. Int. J. Biochem. Cell Biol. 38, 1662–1677.

Joshi, V., and Fernie, A.R. (2017). Citrulline metabolism in plants. Amino acids 49, 1543–1559.

Klareskog, L., Lundberg, K., and Malmström, V. (2013). Autoimmunity in rheumatoid arthritis: citrulline immunity and beyond. Adv. Immunol. 118, 129–158.

Ludidi, N., and Gehring, C. (2003). Identification of a novel protein with guanylyl cyclase activity in Arabidopsis thaliana. J. Biol. Chem. 278, 6490–6494.

Lundberg, K., Wegner, N., Yucel-Lindberg, T., and Venables, P.J. (2010). Periodontitis in RA-the citrullinated enolase connection. Nat. Rev. Rheum. 6, 727–730.

Marondedze, C., Groen, A.J., Thomas, L., Lilley, K.S., and Gehring, C. (2016). A quantitative phosphoproteome analysis of cGMP-dependent cellular responses in Arabidopsis thaliana. Mol. Plant 9, 621–623.

Marondedze, C., Turek, I., Parrott, B., Thomas, L., Jankovic, B., Lilley, K.S., and Gehring, C. (2013). Structural and functional characteristics of cGMP-dependent methionine oxidation in Arabidopsis thaliana proteins. Cell Commun. Signal 11, 1.

Marondedze, C., Wong, A., Groen, A., Serrano, N., Jankovic, B., Lilley, K., Gehring, C., and Thomas, L. (2015). Exploring the Arabidopsis proteome: influence of protein solubilization buffers on proteome coverage. Int. J. Mol. Sci. 16, 857–870.

Pettersen, E.F., Goddard, T.D., Huang, C.C., Couch, G.S., Greenblatt, D.M., Meng, E.C., and Ferrin, T.E. (2004). UCSF Chimera--a visualization system for exploratory research and analysis. J. Comput. Chem. 25, 1605–1612.

Piazza, A., Zimaro, T., Garavaglia, B.S., Ficarra, F.A., Thomas, L., Marondedze, C., Feil, R., Lunn, J.E., Gehring, C., Ottado, J., et al. (2015). The dual nature of trehalose in citrus canker disease: a virulence factor for Xanthomonas citri subsp. citri and a trigger for plant defence responses. J. Exp. Bot. 66, 2795–2811.

Rogers, G.E. (1962). Occurrence of citrulline in proteins. Nature 194, 1149–1151.

Rogers, G.E., and Simmonds, D.H. (1958). Content of citrulline and other amino-acids in a protein of hair follicles. Nature 182, 186–187.

Sali, A., and Blundell, T.L. (1993). Comparative protein modelling by satisfaction of spatial restraints. J. Mol Biol. 234, 779–815.

Smith, D.G., and Young, E.G. (1955). The combined amino acids in several species of marine algae. J. Biol Chem. 217, 845–853.

Tarcsa, E., Marekov, L.N., Mei, G., Melino, G., Lee, S.C., and Steinert, P.M. (1996). Protein unfolding by peptidylarginine deiminase. Substrate specificity and structural relationships of the natural substrates trichohyalin and filaggrin. J. Biol Chem. 271, 30709–30716.

Thompson, P.R., and Fast, W. (2006). Histone citrullination by protein arginine deiminase: is arginine methylation a green light or a roadblock? ACS Chem. Biol. 1, 433–441.

Towbin, H., Staehelin, T., and Gordon, J. (1979). Electrophoretic transfer of proteins from polyacrylamide gels to nitrocellulose sheets: procedure and some applications. Proc. Natl Acad. Sci. USA 76, 4350–4354.

Trott, O., and Olson, A.J. (2010). AutoDock Vina: improving the speed and accuracy of docking with a new scoring function, efficient optimization, and multithreading. J. Comput. Chem. 31, 455–461.

Turek, I., Marondedze, C., Wheeler, J.I., Gehring, C., and Irving, H.R. (2014). Plant natriuretic peptides induce proteins diagnostic for an adaptive response to stress. Front. Plant Sci. 5, 661.

van Venrooij, W.J., and Pruijn, G.J. (2000). Citrullination: a small change for a protein with great consequences for rheumatoid arthritis. Arthritis Res. 2, 249–251.

Yuzhalin, A.E. (2019). Citrullination in Cancer. Cancer Res. 79, 1274–1284.

Zimmermann, P., Hirsch-Hoffmann, M., Hennig, L., and Gruissem, W. (2004). GENEVESTIGATOR. Arabidopsis microarray database and analysis toolbox. Plant Physiol. 136, 2621–2632.

